# Prevalence of chronic chikungunya and associated risks factors in the French West Indies (La Martinique). A prospective cohort study

**DOI:** 10.1101/588970

**Authors:** Bertolotti Antoine, Thioune Marême, Abel Sylvie, Belrose Gilda, Calmont Isabelle, Césaire Raymond, Cervantes Minerva, Fagour Laurence, Javelle Émilie, Lebris Catherine, Najioullah Fatiha, Pierre-François Sandrine, Rozé Benoît, Vigan Marie, Laouénan Cédric, Cabié André, the Chronic Chikungunya working group of University Medical Center of Martinique

**Affiliations:** Université des Antilles, EA 4537, Fort-de-France, France; INSERM, CIC1424, CHU de Martinique, Fort-de-France, France; CHU de Martinique, service de maladies infectieuses et tropicales, Fort-de-France, France; CHU de Martinique, Centre de ressource biologique de la Martinique, Fort-de-France, France; CHU de Martinique, laboratoire de virologie, Fort-de-France, France; INSERM, IAME, UMR 1137; Université Paris Diderot, Paris, France; INSERM, CIC-EC 1425, Hôpital Bichat, Paris, France; Hôpital d’instruction des Armées Laveran, service de pathologie infectieuse et tropicale, Marseille, France; Aix Marseille Université, Institut de Recherche pour le Développement (IRD); Assistance Publique-Hôpitaux de Marseille, Microbes Vecteurs Infections Tropicales et Méditerranéennes (VITROME); Institut Hospitalo-Universitaire Méditerranée Infection, Marseille, France; Département d’Épidémiologie, Biostatistique et Recherche clinique, Assistance Publique-Hôpitaux de Paris, Hôpital Bichat, Paris, France

## Abstract

**Background:** The chikungunya virus (CHIKV) is a re-emerging alphavirus that can cause chronic rheumatic musculoskeletal disorders, named chronic chikungunya (CHIKC+), which may be long-term incapacitating. A chikungunya outbreak occurred in 2013 in La Martinique. We constituted the first prospective cohort study of CHIKV-infected subjects in the Caribbean to assess the prevalence of CHIKC+ at 12 months and to search for factors present at the acute stage significantly associated with chronicity.

**Methodology/Principal findings:** A total of 193 patients who tested positive for RT-PCR CHIKV, were submitted to clinical investigations in the acute phase (<21 days), and 3, 6, and 12 months after their inclusion. A total of 167 participants could be classified as either suffering or not from CHIKC+. They were analyzed using logistic regression models. At 12 months, the overall prevalence of CHIKC+ was 52.1% (95%CI: 44.5-59.7). In univariate analysis, age (OR: 1.04; 95% CI: 1.02-1.07; p=0.0003), being male (OR: 0.51; 95%CI: 0.27-0.98; p=0.04), headache (OR: 1.90; 95%CI: 1.02-3.56; p=0.04), vertigo (OR: 2.06; 95%CI: 1.05-4.03; p=0.04), vomiting (OR: 2.51; 95%CI: 1.07-5.87; p = 0.03), urea (OR: 1.33; 95%CI: 1.05-1.70; p=0.02) were associated with CHIKC+. In final multivariate logistic regression models for 167 participants, predictors of CHIKC+ were age (OR 1.06; 95%CI: 1.03-1.08; p<0.0001), male sex (OR: 0.40; 95%CI: 0.19-0.84; p=0.015), vertigo (OR: 2.46; 95%CI: 1.16-5.20; p=0.019), hypotension (OR 4.72; 95% -CI: 1.19-18.79; p=0.028), recoloration time >3 seconds (OR: 3.79; 95%-CI: 1.01-14.25).

**Conclusions:** This cohort study in La Martinique confirms that CHIKC+ is a frequent complication of acute chikungunya disease. Analysis emphasized the importance of age and female sex for CHIKC occurrence, and pointed out the potential aggravating role of dehydration at the acute phase. Early and adequate hydration could reduce the risk chronic chikungunya disorders.

**Author Summary:** Chikungunya is a mosquito-borne virus found in tropical countries that has been re-emerging in the last decade. It has caused major epidemics in recent years, such as in Reunion Island and in Southeast Asia. Nearly 2.5 billion people around the world are at risk of contracting the virus. During the acute phase of the illness, patients experience a flu-like syndrome with fever, headache, myalgia, rash, and severe arthralgia. These symptoms can persist for several months in some patients, and can lead to significant functional disability. During the 2013 epidemic in Martinique, we followed nearly 200 patients who had contracted chikungunya. More than half of the patients had a chronic form of the disease—mainly women over 50 years of age. Our statistical analyses indicate that poor hydration during the acute phase may be a risk factor for developing chronic rheumatism. Therefore, in the context of a chikungunya epidemic, it seems important to encourage patients to drink plenty of fluids as soon as the first symptoms appear.

## Introduction

Chikungunya virus (CHIKV), “that which bends up” in the Makonde dialect [1], is a re-emerging alphavirus transmitted to human by *Aedes* mosquitoes, which has caused massive epidemics in Africa [2-4]; in the Indian Ocean islands [5-7]; in Southeast Asia [8,9]; and, in 2007, in Italy, which was the first European outbreak [10,11]. CHIKV is known to target human epithelial and endothelial cells, fibroblasts and macrophages [12-14], and human muscle satellite cells [15]. The virus is also suspected of neurotropism [16,17], regarding its neurological complications [18-21].

The disease typically consists of an acute illness like dengue fever, characterized by abrupt onset of a high-grade fever, followed by constitutional symptoms, poly-arthritis, musculoskeletal pain, headache, skin involvement [22], and sometimes severe manifestations like encephalopathy [20,23] acute hepatitis [24,25], myocarditis [26-28], and multi-organ failure. Usually, the acute phase lasts on average 5 to 10 days (viremic period), and resolves within a few weeks. In May 2015, the World Health Organization [29] defined a person with chronic chikungunya as a “person with previous clinical diagnosis of chikungunya after 12 weeks of the onset of the symptoms presenting with at least one of the articular manifestations: pain, rigidity, or edema, continuously or recurrently.” Chronic arthralgia post CHIKV-infection (i.e. “chronic chikungunya” or “CHIKC+”) is a frequent complication. A total of 40 to 60% [30-32] of acute chikungunya cases reported a significant long-term impaired quality of life [33-36]. Previous studies from other outbreaks suggested that patients infected with CHIKC+ were significantly older [30,32,37,38], rather females [7,30,38,39], had severe initial rheumatic symptoms [38,40,41], and had high CHIK-specific IgG titers [37,40]. Notably, initial predictive factors of CHIKC were robustly but retrospectively investigated by Gerardin *et al.* [40] in a cohort of 346 patients infected during the 2005-2006 epidemic in La Reunion. La Martinique, is a French overseas department of nearly 400 000 inhabitants, located in the French West Indies. The first autochthonous cases of chikungunya were described in the French West Indies in November 2013 (December 2013 in La Martinique) [42]. At the end of the outbreak (approximately in January 2015), the number of affected people was estimated 145,000 (36% of the population) [43]. This study had two objectives: first, to define the prevalence of CHIKC+ at 12 months of follow-up, and secondly, to investigate potential risk factors of CHIKC+.

## Methods

Ethical clearance was obtained by ANSM (n°IDRCB 2010-A00282-37) and by the committee for the protection of individuals. Written and signed informed consent of all subjects was obtained.

The Department of Infectious Diseases of the Hospital of Fort de France started the “DAG” Study in June 2010. This is a descriptive and prognostic study of dengue fever in the French West Indies and French Guiana, based on a hospital cohort of children and adults with suspected dengue fever. The main objective of this study was to define the predictive factors of severe dengue disease. Following the CHIKV introduction in La Martinique, inclusion criteria of the DAG study were extended also to chikungunya cases, with a monitoring adjustment (including a wide description of rheumatic disorders specific to chikungunya disease). In May 2014, the “DAG” study became the “DAG2” study, with a cohort composed of two groups: the dengue group and the chikungunya group. This study is registered on clinicaltrials.gov (NCT01099852). The DAG2 study is still ongoing (the duration of the follow-up is 36 months), and this analysis was conducted at 12 months’ follow-up.

At the beginning of 2016, the DAG2 cohort was extended to other arbovirosis (including the Zika virus), which has been renamed “the CARBO cohort”.

This study was proposed to the departments of the Hospital of Fort de France which were likely to receive patients infected by the CHIKV (the emergency department and department of infectious diseases). Therefore, the cohort of patients was ambulatory and reflected so the global population from Martinique affected during the epidemic.

### Study population

For this study, patient inclusion was conducted from December 19, 2013, to December 4, 2014. Inclusion criteria were: age > 16 years old, ability to accept and sign informed consent, positive blood CHIKV qRT-PCR, and onset of symptoms ≤7 days.

### Data collection and follow-up

The DAG questionnaire was adapted to the purpose of our study. Data at the acute stage were collected at the initial visit, and if timely possible at day 3 after the onset of symptoms, between day 5 and 7, and between day 8 and 10 including : sociodemographic data, comorbidities and clinical characteristics at baseline (headache, arthralgia myalgia, fever, rash, fatigue, etc.), impact on quality of life measured by the EuroQol 5 Dimensions (EQ5D) assessment and treatments used. For patients enrolled from July 2014 (after adjustment of clinical files to the DAG2 study) an accurate description of affected joints (location, swelling, stiffness, arthritis, enthesitis trouble, joint pain, etc.), and an assessment of peripheral neuropathy as measured by the DN4 questionnaire were also included. At 3 months (M3), patients were examined by a physician of the team to identify CHIKC+ cases. At 6 months (M6) and 12 months (M12), patients were interviewed by telephone using the same questionnaire made up of closed questions to monitor persistent symptoms notably joint pains, the impact on the quality of life (measured by the “EQ5D” assessment), and the treatments received. A subject was definitively classified as suffering from CHIKC+ if the answer was NO at any time of the follow-up from M3 to the question: “Do you feel completely recovered from CHIKV-infection?”. Therefore a clinical examination was proposed to describe and to manage remaining disorders. Only patients who felt completely recovered from chikungunya at all the visits completed from M3 were categorized as CHIKC- (no chronic chikungunya disease). The global clinical assessment of CHIKC+ subjects, were estimated using the “Multi-Dimensional Health Assessment Questionnaire” (MDHAQ). Usually, patients with chikungunya disease describe a fluctuation of their symptoms (continuously or recurrently) from day to day. Patients with relapsing pain were defined as having “disorders present at one and/or two time points without recovering” and patients with lingering pain were defined as having “disorders present at the three time points: M3, M6, and M12.”

The 12-months-period of follow-up enabled to widely describe the chronic chikungunya disorders and to capture for each patient the worst time-point with the heaviest burden (M3, M6, or M12).

### Laboratory tests

Only CHIKV-viremic patients were recruited. Blood cells counts, biochemical tests : notably C-reactive protein (CRP), ionogram, urea, creatinine and hepatic transaminases, and a biological database (serotheque, plasmatheque, cellulotheque, DNAtheque, RNA theque) were performed at the time of positive plasmatic CHIKV qRT-PCR. However, the viral load was not collected in this study.

### Data Analyses and Statistical Tests

Results were expressed as mean and standard error or frequencies (percentages) as appropriate. Univariate and multivariate logistic regression analyses were performed to evaluate factors associated with CHIKC+. The factors associated with CHIKC+ with a P value <0.25 in the univariate analysis, were included in a multivariate model and then selected using a backward stepwise strategy with a P value <0.05. Odds ratios and 95% confidence intervals were estimated. Some data significant in univariate analyses were incomplete or partially completed, but equally in all groups. Missing values were imputed by fully conditional specification methods with 15 imputed datasets.

For the subgroup who had clinical data on joints involvement, comparisons were between CHIKC- and CHIKC+ were performed using an univariate analysis. Due to low effectives and risk of overfitting, multivariate analysis was not conducted in this subpopulation.

All statistical analyses were performed with SAS® 9.4 (SAS Institute, Cary, NC, USA).

## Results

A total of 247 were eligible and consented to participate, of whom 193 were enrolled (54 exclusions: 51 had negative qRT-PCR, 2 were not tested for CHIKV qRT-PCR, and one was over the 7 days delay since the onset of symptoms for inclusion). The flow-chart of the study is displayed in Fig 1.

**Fig 1.**
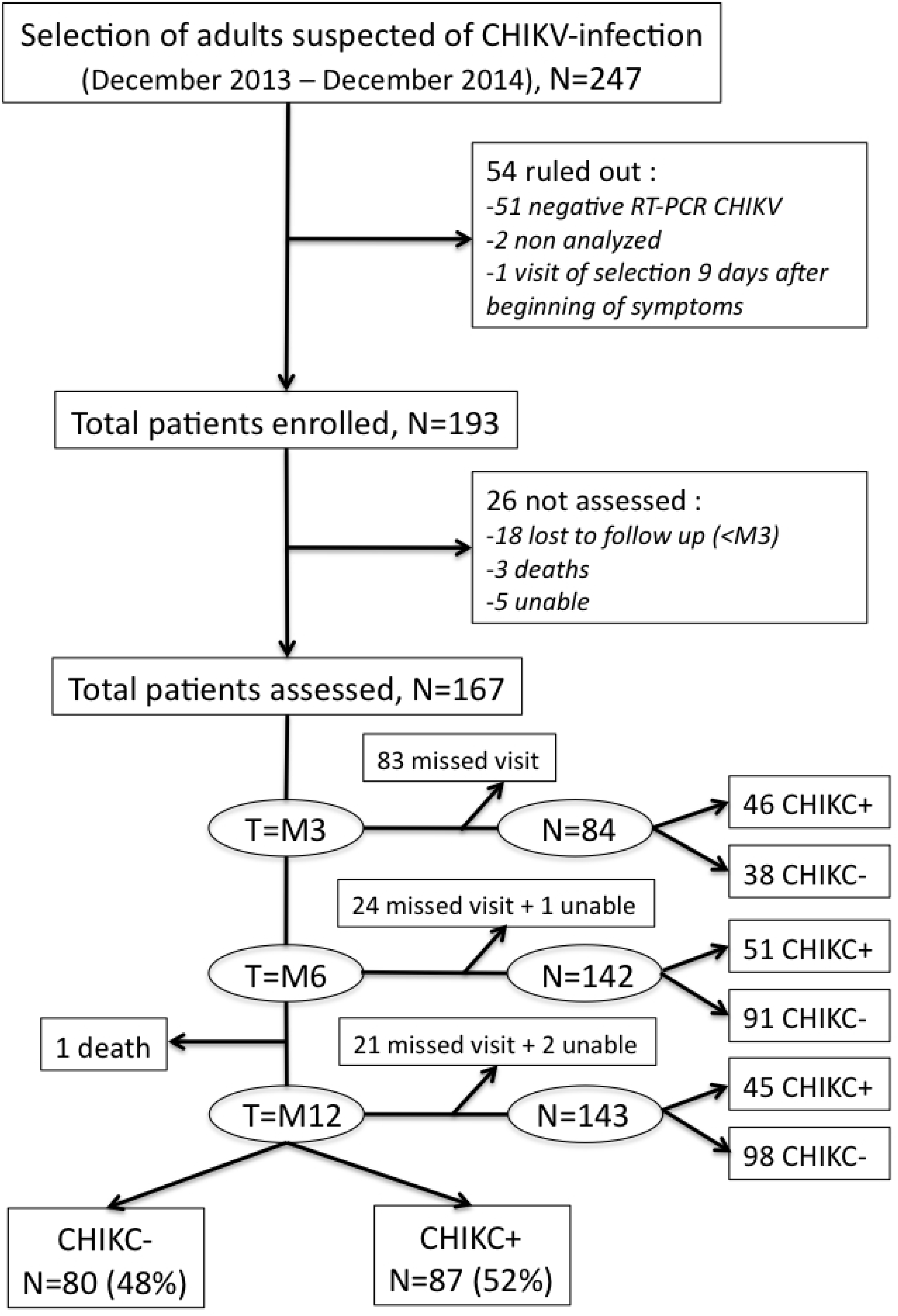
Diagram of the study population. “unable”: patients unable to answer the question “do you feel completely recovered from CHIKV-infection?” “missed visit”: not present at a specific time point. “lost to follow-up”: patients who missed all visits from M3. “death”: dead patients. “CHIKC+”: patients with chronic chikungunya disease. “CHIKC–”: patients chronic chikungunya disease-free.

### Overall prevalence of CHIKC+ at 12 months

Among 193 patients, 21 were lost-to follow-up (i.e. reached neither M3 nor M6 nor M12), of whom 3 had died before M3 (causes of death were: adenocarcinoma of the prostate, multi-organ failure at the acute CHIKV-infection, and unknown cause). Between M6 and M12 one patient died of myeloma. Five subjects have remained with undetermined CHIKC-status during all their follow-up and were excluded from the analysis.

A total of 167 patients could be classified as either CHIKC- or CHIKC+. Close to half patients missed the M3 visit since only 46 CHIKC+ patients and 38 CHIKC-patients were counted at that time. However, more were evaluated at M6 and M12, with respectively 142 (85%; 51 CHIKC+ and 91 CHIKC-) and 143 (86%; 45 CHIKC+ and 98 CHIKC-) patients classified. A total of 87 patients were CHIKC+ among the 167 participants (80 were CHIKC-), giving an overall prevalence of 52.10% (CI 95% 44.5-59.7). Ten CHIKC+ subjects attended at only one visit, so their CHIKC profile (“relapsing” or “lingering”) couldn’t be assessed.. Most CHIKC+ patients had relapsing CHIKC+ 83.12% (64 out of 77), versus 16.88% (13 out of 77) with lingering CHIKC+.

### Clinical data at the acute stage of the global population

All the clinical and biological characteristics at the onset of disease are displayed in the S1 and S2 Tables.

The ratio of men to women was 0.56 (59 males and 106 females). Of the 167 patients included (age range from 20 to 91 years), 53.9% were aged over 50 years, with a mean (±SD) age of 51.35 (±16.10) years. Amerindian (n=65, 38.92%) and Sub-Saharan Africa origins (n=50, 29.94%) were mainly represented. Cardiovascular and rheumatologic diseases were the most common pre-existing comorbidities: 36 (21.56%) high blood pressure, 28 (16.77%) healed fractures, 14 (8.38%) osteoarthritis, 13 (7.78%) diabetes, 13 (7.78%) dyslipidemia, 11 (6.59%) rheumatologic inflammatory diseases, and 9 (5.39%) cancer.

The most frequently recorded clinical signs at the acute phase were fever (95.21%), intensity of pain > 4/10 on the visual analog scale (83.83%), headache (59.28%), myalgia (56.29%), and vertigo (31.74%); digestive symptoms were less frequent (diarrhea, vomiting, and abdominal pain).

For the subpopulation with data on joints involvement at the disease onset (n=73 with 37 CHIKC- and 36 CHIKC+ at the end),, the majority had articular disorders (69 patients, 94.52%) with joint pain (90.41%), swelling (42.46%), stiffness (27.40%), arthritis (20.55%), tenosynovitis (13.70%), and enthesitis (10.96%). The joints most affected were the ankles (68.49%), the wrists (58.90%), the distal interphalangeal joints in hands (36.97%), the knees (50.68%), and the proximal interphalangeal in hands (45.20%).

### Identification of risk factors for chronic outcome (CHIKC+)

To search for the initial predictive factors for CHIKC+, characteristics at disease onset in the two groups (CHIKC- and CHIKC+) were compared, and detailed in S1 and S2 Tables. Significant results are listed in table 1. Patients CHIKC+ were mainly women aged over 50 years (Fig 2). In this study, 58% (61/106) of the women developed CHIKC, versus 41% (24/59) of the men (gender data available for 165 patients). In univariate analysis (Table 1), the probability of being CHIKC+ increased with age (OR 1.04, 95% CI, 1.02-1.07, p=0.0003); female gender (OR 0.51, 95% CI, 0.27-0.97, p=0.04); and some clinical signs at the disease onset like headache (OR 1.90, 95% CI, 1.02-3.56 p=0.04), vertigo (OR 2.06, 95% CI, 1.05-4.03, p=0.04), vomiting (OR 2.51, 95% CI, 1.07-5.87, p=0.03), and dyspnea (OR 2.12, 95% CI, 1.02-4.42, p=0.04). In the subpopulation better characterized in terms of joint involvement at the acute stage at least one enthesitis (OR 8.69, 95%CI, 1.01-74.72, p=0.05) was significantly associated with CHIKC+ in the univariate analyses, while at least one tenosynovitis (OR 5.0, 95% CI, 0.98-25.45, p=0.05) didn’t reach the statistical significance (Table 2). Biologically, the increase of urea (OR 1.33, 95% CI, 1.0.5-1.70, p=0.02) was significantly associated with evolving CHIKC+. No comorbidity (cardiovascular or rheumatologic diseases) was found to be a risk factor for being CHIKC+ (S1 Table and Table 1). Impact on quality of life as measured by the EQ5D score at the acute phase was not significantly different between patients in the CHIKC- and CHIKC+ groups (S1 Table).

**Table 1.** Significant characteristics of CHIKC+ and CHIKC- adult at disease onset. Univariate and multivariate analysis. DAG-2 study 2014-2016, La Martinique (N=167).

| Variables | OR [IC95%] | p-value | aOR [IC95%] | p-value* |
| --- | --- | --- | --- | --- |
| Age (year) | 1.04 [1.02 – 1.06] | **0.0003** | 1.06 [1.03 – 1.08] | **<0.0001** |
| Male | 0.51 [0.27 – 0.98] | **0.04** | 0.40 [0.19 – 0.84] | **0.015** |
| Headache | 1.90 [1.02 – 3.56] | **0.04** |  |  |
| Vertigo | 2.06 [1.05 – 4.03] | **0.04** | 2.46 [1.16 – 5.20] | **0.019** |
| Vomiting | 2.51 [1.07 – 5.87] | **0.03** |  |  |
| Nausea | 2.35 [0.91 – 6.05] | 0.08 |  |  |
| Dyspnea | 2.12 [1.02 – 4.42] | **0.04** |  |  |
| Confusional syndrome | 0.22 [0.02 – 2.02] | 0.18 |  |  |
| Low blood pressure (SBP < 90 mmhg and DBP < 60 mmhg) | 2.26 [0.68 – 7.50] | 0.18 | 4.72 [1.19 – 18.79] | **0.028** |
| Recoloration time > 3 sec | 2.42 [0.76 – 7.75] | 0.14 | 3.79 [1.01 – 14.25] | **0.049** |
| Pulse (beats per minute) | 0.99 [0.97 – 1.01] | 0.25 |  |  |
| Score scale EQ5D in the acute phase | 0.99 [0.97 – 1.00] | 0.12 |  |  |
| Thrombopathy or chronic thrombopenia | 5.85 [0.70 – 49.67] | 0.11 |  |  |
| Hemoglobinopathy | 4.82 [0.55 – 42.14] | 0.16 |  |  |
| Antihistaminic | 1.50 [0.76 – 2.94] | 0.24 |  |  |
| Non-steroidal anti-inflammatory drugs | 1.76 [0.87 – 3.54] | 0.11 |  |  |
| Red blood cells (G/L) | 0.67 [0.38 – 1.17] | 0.16 |  |  |
| Hemoglobin (G/dl) | 0.88 [0.72 – 1.07] | 0.19 |  |  |
| Eosinophils (G/L) | 0.07 [0.001 – 3.05] | 0.07 |  |  |
| Urea (mmol/L) | 1.33 [1.04 – 1.69] | **0.02** |  |  |

OR : odd ratio; aOR : adjusted odd ratio; \*wald test

**Table 2.** Joints involvement at the CHIKV acute infection in a subpopulation of the DAG-2 study 2014-2016, La Martinique (N=73) : univariate comparison between CHIKC+ and CHIKC- adult.

| Variables | N | CHIKC- (%)<br>(N=37) | CHIK+ (%)<br>(N=36) | OR [IC95%] | P-value* |
| --- | --- | --- | --- | --- | --- |

| Sites |  |  |  |  |  |
| --- | --- | --- | --- | --- | --- |
| Ankles | 73 | 26 (70.3) | 24 (66.7) | 0.85 [0.32 – 2.27] | 0.7 |
| Wrists | 73 | 22 (59.5) | 21 (58.3) | 0.96 [0.38 – 2.42] | 0.9 |
| Distal interphalangeal joint (hand) | 73 | 10 (27.0) | 17 (47.22) | 2.41 [0.91 – 6.42] | 0.08 |
| Knees | 73 | 18 (48.7) | 19 (52.8) | 1.18 [0.47 – 2.96] | 0.7 |
| Shoulder | 73 | 17 (46.0) | 17 (47.2) | 1.05 [0.42 – 2.64] | 0.9 |
| Proximal interphalangeal joint (hand) | 73 | 16 (43.2) | 17 (47.2) | 1.17 [0.47 – 2.95] | 0.7 |
| Metacarpophalangeal joint | 73 | 11 (29.7) | 14 (38.9) | 1.50 [0.57 – 3.98] | 0.4 |
| Elbow | 73 | 10 (27.0) | 10 (27.8) | 1.04 [0.37 – 2.91] | 0.9 |
| Metatarsophalangeal joint | 73 | 9 (24.3) | 9 (25.0) | 1.04 [0.36 – 3.01] | 0.9 |
| Foot | 73 | 9 (24.3) | 8 (22.2) | 0.89 [0.30 – 2.64] | 0.8 |
| Hip | 73 | 7 (18.9) | 8 (22.2) | 1.22 [0.39 – 3.82] | 0.7 |
| Hand | 73 | 5 (13.5) | 6 (16.7) | 1.28 [0.35 – 4.64] | 0.7 |
| Ribs | 73 | 0 | 1 (2.78) | - |  |
| Back | 73 | 5 (13.51) | 7 (19.44) | 1.55 [0.44 – 5.41] | 0.5 |
| Neck | 73 | 10 (27.03) | 13 (36.11) | 1.53 [0.57 – 4.13] | 0.4 |
| Rachis | 73 | 16 (43.24) | 10 (27.78) | 0.51 [0.19 – 1.34] | 0.2 |
| Sternum | 73 | 2 (5.41) | 5 (13.89) | 2.82 [0.51 – 15.60] | 0.2 |
| Proximal interphalangeal joint (foot) | 73 | 5 (16.5) | 5 (13.9) | 1.03 [0.27 – 3.92] | 1.0 |
| Distal interphalangeal joint (foot) | 73 | 4 (10.8) | 5 (13.9) | 1.33 [0.33 – 5.41] | 0.7 |
| Articular disorders |  |  |  |  |  |
| Joint pain | 73 | 33 (89.2) | 33 (91.7) | 1.33 [0.28 – 6.43] | 0.7 |
| Joint swelling without arthritis | 73 | 17 (46.0) | 14 (38.9) | 0.75 [0.30 – 1.90] | 0.5 |
| Joint stiffness | 73 | 7 (18.9) | 13 (36.1) | 2.42 [0.83 – 7.04] | 0.1 |
| Arthritis | 73 | 7 (18.9) | 8 (22.2) | 1.22 [0.39 – 3.82] | 0.7 |
| Asymmetry joint achievement | 73 | 7 (18.9) | 5 (13.9) | 0.69 [0.20 – 2.42] | 0.6 |
| **Tenosynovitis** | **73** | **2 (5.4)** | **8 (22.2)** | **5.00 [0.98 -25.45]** | **0.05** |
| **Enthesitis** | **73** | **1 (2.7)** | **7 (19.4)** | **8.69 [1.01 – 74.72]** | **0.049** |
| Arthritis with synovitis | 73 | 1 (2.7) | 0 | - | - |
| Periostitis | 73 | 0 | 0 | - | - |

\*wald test

**Fig 2.**
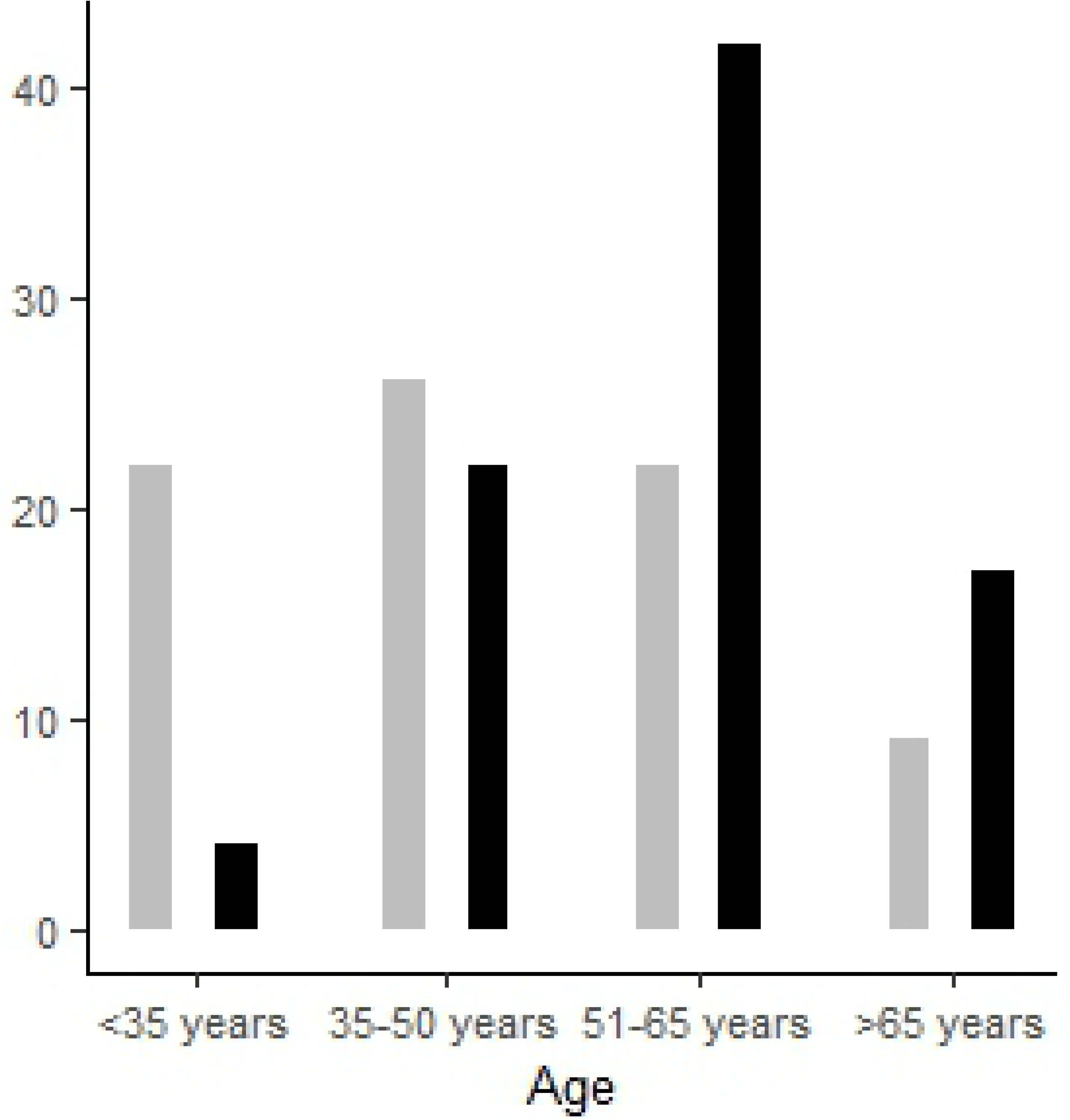
Age distribution in the DAG-2 study 2014-2016, La Martinique. In Grey, for 80 CHIKC-. In Black, for 87 CHIKC+.

In the multivariate logistic regression model presented in Table 1, the predictors of CHIKC+ were the age (OR 1.06, 95% CI, 1.03-1.0.8, p<0.0001), and vertigo (OR 2.46, 95% CI, 1.16-5.20, p=0.02), low blood pressure (OR 4.72, 95% CI, 1.19-18.79, p=0.03) or recoloration time > 3 seconds (OR 3.79, 95%CI, 1.01-14.25, p=0.049) at the time of acute infection. Male gender (OR 0.40, 95% CI, 0.19-0.84, p=0.02) was protective.

Relapsing-remitting joint symptoms occurred in 64 of 77 answering chronic patients (83%), while only 17% (13/77) had a continuous course. A total of 54 of 87 CHIKC+ patients were clinically evaluated at the chronic phase (Table 3), with joint pain (87%), stiffness (33.3%), and swelling (22%) being the most common chronic signs. Enthesitis (13%), arthritis (7.4%), and tenosynovitis (5.6%) were observed less frequently. Ankles (31.5%) and knees (29.6%) were the most affected joints, while the proximal interphalangeal joints in hands (24.1%), the wrists (22.2%), and the shoulders (22.2%) were less frequently involved. A quarter of patients had peripheral neuropathy (27.7%), a mean MDHAQ score of 10 out of 30, and a decrease in their quality of life (mean score EQ5D = 51.56/100).

**Table 3.**
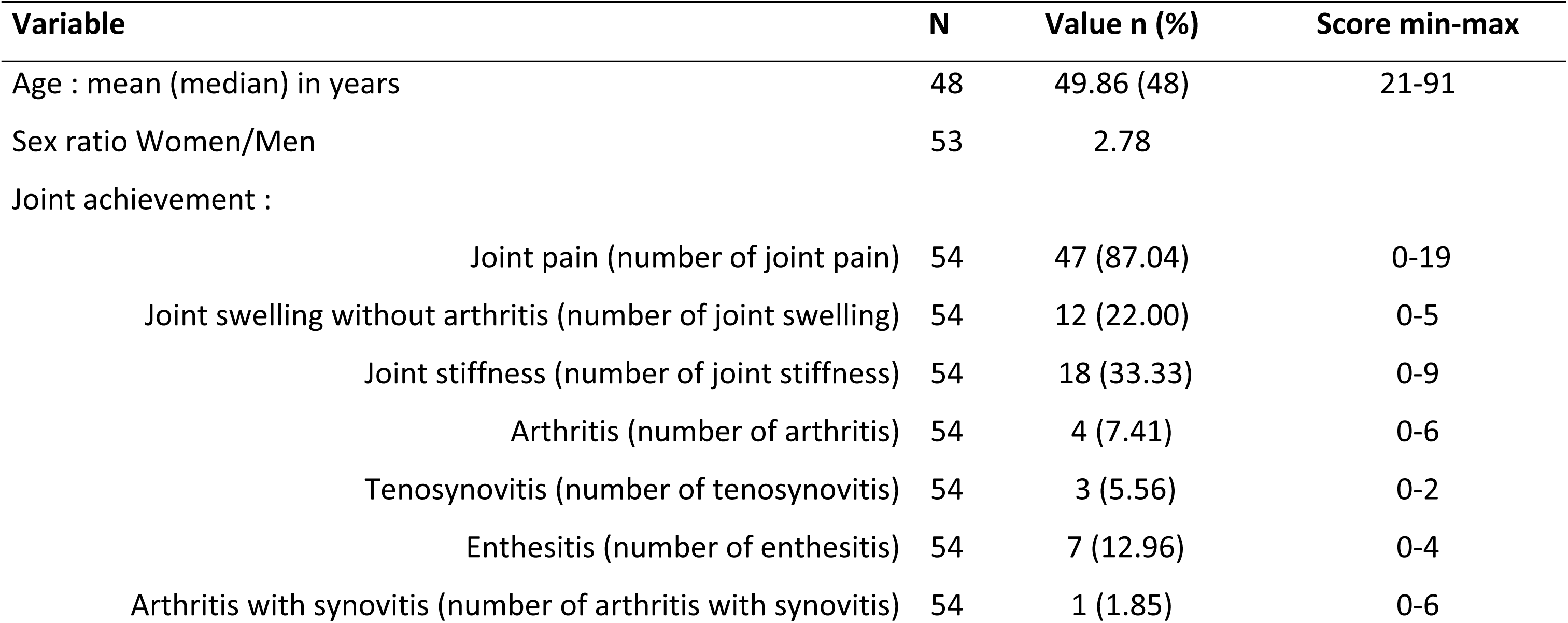
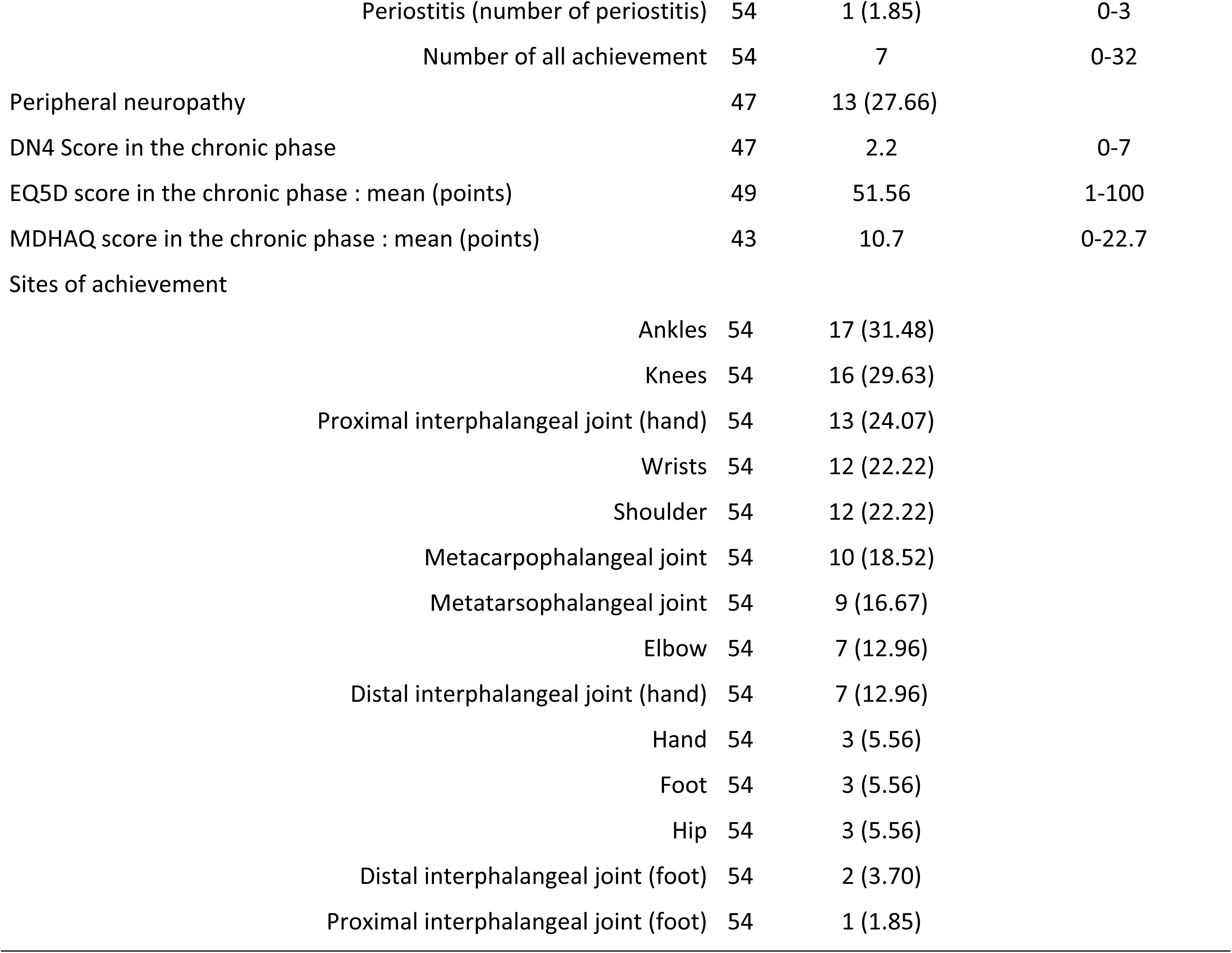
Clinical profile of CHIKC+ patients in the chronic phase (N=54).

## Discussion

This is the first prospective cohort study of CHIKV-infected cases in La Martinique during the 2013-outbreak [43]. The global prevalence of chronic chikungunya was 52.10% (CI 95% 44.5-59.7), confirming that it is a frequent evolution after the acute infection as it was previously observed [7,30-32], except in India were the lowest rate was reported [46,47] (possibly due to viral or population specificities).

The age over 50 and the female gender were also assessed as main risk factors for chronic chikungunya disease during previous outbreaks [7,30,32,38-40,48]. Some theories related to the gender have emerged such as the role of the menstrual cycle [49] and the ovulation influencing the production of pro-inflammatory cytokines by monocytes [49,50], or the role of estradiol in the increase of antibodies production [51,52] but nothing has been proved.

Cartilage is essentially composed of water [53]; therefore, dehydration may result in its degradation by causing cracks and increasing fragility [54,55]. In our study, some initial factors significantly associated with CHIKC+ in univariate and multivariate analyses -notably vertigo, headache, vomiting, nausea, hypotension, recoloration time > 3 seconds, and increased urea- may be linked with dehydration. Moreover, intravenous rehydration was found less frequently implemented at the acute stage in the group CHIKC+ than in the group CHIKC- (*p=0.05*). This raises the hypothesis of a relationship between dehydration and the risk of CHIKC+ development.

Recent data by Boettcher *et al.* [56] suggest that after dehydration, the transient structure and mechanical properties of articular cartilage decreases the risk of irreversible material failure, despite cartilage’s poor regenerative ability. Other research by Sahni *et al.* [57], using mice, focused on the contribution of prolonged dehydration on osseous tissue and calcium phosphate metabolism. They demonstrated that prolonged water deficiency would entail a decreased bone mass, with an increase of rate of bone resorption and mineralization defect. In the same way, calcium metabolism disorders stimulate parathyroid hormone (PTH) secretion and activation of bone resorption by osteoclast. In our study, however, calcemia didn’t differ between CHIKC+ and CHIKC- patients at the acute stage, but the sample was small (calcium rate available for 61 patients only) and it was a one-off measure that can’t reflect metabolic disorders over the medium to long term. Moreover hydration could help dilute pro-inflammatory cells and cytokines recruited in joints and enhance anti-inflammatory response as the water on fire.

It has been demonstrated that CHIKV is able to damage cartilage [58] and synovial tissue [59]. In this context, the hypothesis would be that “at disease onset of chikungunya, patients suffering in addition from dehydration could have a lower resilience for developing damaged cartilage. Joints injury could be more severe with a slower recovery and with a higher risk of chronicity.” Furthermore, it has been reported that the intensity of symptoms at presentation was predictive of CHIKC+ [40]. Dehydration could also participate in the increase in viremia through serum concentration. Higher CHIKV plasma viral loads have been reported in CHIKC+ patients [59,60], but we haven’t investigated this parameter in our study.

Contrary to other authors, we didn’t find pre-existing diseases more likely associated with CHIKC+, neither rheumatic disorders [48] such as osteoarthritis [39,61] and rheumatoid arthritis [62-66], nor cardiovascular diseases including hypertension [67] and diabetes [32].

Inflammatory and autoimmune processes in chikungunya disease at the acute and chronic satges have been described in recent studies [59,68]. Several of them have identified the positive relationship at disease onset between CRP level [32,59], pro-inflammatory cytokines [59,68], and severity at presentation [40,41,61]; however, no autoimmune markers for CHIKV were suggested in observational studies [32]. In other studies, biomarkers associated with CHIKC+ were IL-6 secretion [60] at disease onset and GM-CSF [59,60]. In the La Reunion Island cohort, the proper role of specific IgG titers for predicting CHIKC+ has clearly been observed and positively correlated with age, female sex, and severity of initial rheumatic symptoms [40]. In this study, the CRP level at disease onset wasn’t associated with CHIKC+.

The main clinical characteristics of chronic patients were consistent with previous descriptions [9,22,36]: polyarthralgia, articular stiffness, swollen joints and arthritis. We also observed enthesitis and tenosynovitis [36,69,70,71]. Relapsing-remitting arthralgia were reported by the majority of patients[40]. We found tenosynovitis, or enthesitis occurrence at the acute stage of chikungunya (at least one of these) associated with the development of CHIKC+. These both conditions highly specific to CHIKV have never been yet identified as risk factors for CHIKC+ [39,72-74].

The impact of CHIKC+ on activities of daily living and quality of life was also investigated [33-36,40]. Patients suffering from CHIKC+ must have a multidisciplinary medical care plan [75,76]. A general practitioner must follow up the patient’s condition long term, often in cooperation with a rheumatologist or the rehabilitation physician [77]. Medical treatments employed for patients are analgesics, anti-inflammatory drugs, corticosteroid, and sometimes anticonvulsant and immunosuppressive therapy in severe cases [76,78,79]. Treatment may be combined with physiotherapy, physical therapy, sychiatric and psychologic [80-82]. Chikungunya disease and specially CHIKC+ with relapsing evolution can induce anxiety, stress, and sometimes a real nervous breakdown [81,82]. Long-term chikungunya disabilities are costly at individual and collective scales. The economic impact of the CHIKV 2005-2006 outbreak in La Reunion was estimated 51.63 million US dollars [83].

One of the strengths of our study was the early inclusion and the prospective follow-up at the epicenter of the epidemic. Moreover, patients were examined physically by a clinician investigator at disease onset and at M3 to monitor clinical manifestations. This objective assessment of symptoms has probably prevented an overestimation of symptoms by the patients; however, some data are subjective as the joint pain assessment.

This study, however, has some limitations. First, the phone interview at M6 and M12 could have contributed to a declarative bias. On the other hand, phone calls limit the rate of lost-to-follow-up.

The plasma viral load in CHIKC+ and CHIKC- patients might be a track to explore using the biobank of the cohort, together with other immunological and virological investigations.

To conclude, this first prospective study during the CHIKV outbreak in La Martinique describes the course of the chikungunya disease in the local population, and helps in estimating the burden at one year. Our findings confirm that chronic stage concern more than half symptomatic infected-people with age and female gender as major risk factors. Some predictors of chronicity identified at the acute stage have introduce for the first time the hypothesis of the dehydration as a compounding factor in persisting rheumatic disorders which opens up new perspectives for prevention and research in CHIKV pathways.

## Supporting information

**S1 Table.** Clinical characteristics of CHIKC+ and CHIKC- adult at disease onset. DAG-2 study 2014-2016, La Martinique (N=167). sd : standard deviation; * wald test

**S2 Table.** Results of biological tests in CHIKC+ and CHIKC- adult at time of chikungunya acute infection. Univariate analysis, DAG-2 study 2014-2016, La Martinique (N=167). *Wald test

**S3 Checklist:** STROBE Checklist

## Acknowledgments

We would like to thank the english language editing from Elsevier for editorial assistance and the Chronic Chikungunya working group of University Medical Center of Martinique (Bally Jacques, Brouste Yannick, Brunier Lauren, Godaert-Simon Ludivine, Hochedez Patrick, Jean-Marie Janick, Komla-Souhka Isabelle, Molkkar Sabine, Pelonde-Erimée Véronique, Perreau Caroline, René-Corail Patrick, Signate Aissatou, Trois-Gros Odile, Ursulet Gilbert, Valentino Ruddy).

The protocol was prepared with the help of the INSERM Research and Action Targeting Emerging Infectious Disease (REACTing) network.

## Author Contributions

**Conceptualization:** Andre Cabie

**Data curation:** Calmont Isabelle

**Formal analysis:** Antoine Bertolotti, Mareme Thioune

**Investigation:** Abel Sylvie, Belrose Gilda, Césaire Raymond, Fagour Laurence, Lebris Catherine, Najioullah Fatiha, Pierre-François Sandrine, Rozé Benoît

**Methodology:** Antoine Bertolotti, Cedric Laouenan

**Supervision:** Andre Cabie

**Validation:** Cedric Laouenan, Marie Vigan

**Visualization:** Minerva Cervantes

**Writing ± original draft:** Antoine Bertolotti, Mareme Thioune

**Writing ± review & editing:** Antoine Bertolotti, Mareme Thioune, Andre Cabie, Cedric Laouenan, Minerva Cervantes, Emilie Javelle

